# *In Vitro* Budding and Flotation-based Enrichment of Mitochondria-derived Vesicles for Proteomics from Mammalian Hearts

**DOI:** 10.1101/2024.03.04.583262

**Authors:** Thejaswitha Rajeev, Ela Lepcha, Nidhi Nair, Lariza Ramesh, Areeba Marib, John B Johnson, Dhabaleswar Patra, Ananthalakshmy Sundararaman

## Abstract

Mitochondria-derived vesicles (MDVs) are carriers involved in mitochondrial quality control, yet their biogenesis and trafficking remain poorly understood. We describe an *in vitro* protocol to generate MDVs from animal heart mitochondria. The workflow allows for isolation of distinct MDV subsets and their proteomic analyses. The protocol improves vesicle yield and purity for NTA, TEM, or western blotting-based readouts. This facilitates the identification of novel MDV cargo and regulators of MDV biogenesis in different model organisms.

For complete details on the use and execution of this protocol, please refer to Rajeev T *et al*., ^[1]^.

**Graphical abstract:** 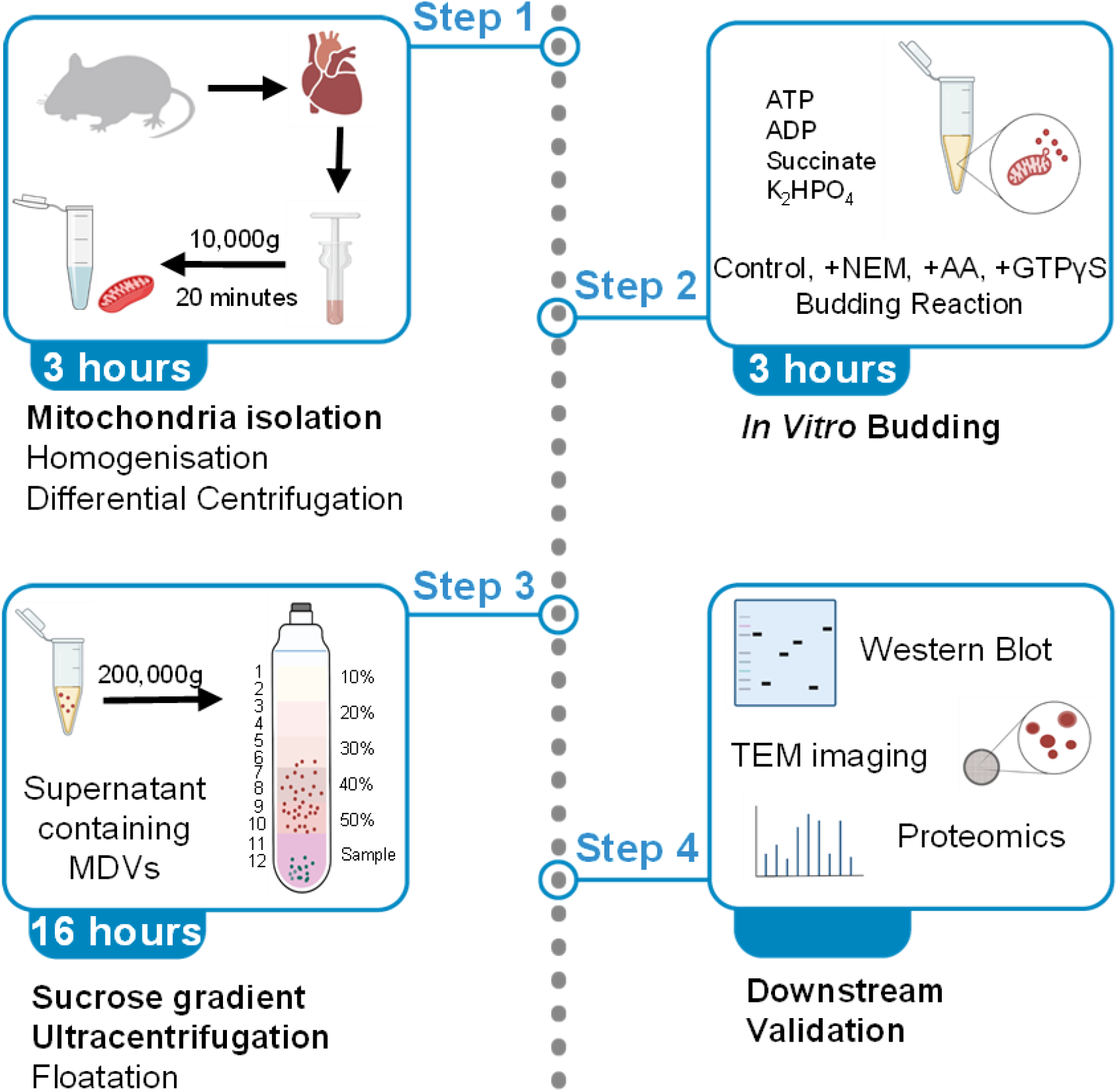

## Before you begin

This protocol describes the isolation of mitochondria from rat heart tissue, followed by an *in vitro* budding to generate mitochondria-derived vesicles (MDVs) from purified mitochondria. The protocol has been adapted and further modified from Soubannier et al.^[2]^ with modifications to optimize mitochondrial yield from cardiac tissue and to improve the separation of budded vesicles on discontinuous sucrose gradients.

Although optimized for rat heart, the protocol has also been successfully applied to rabbit and bovine heart tissues.

As the mitochondrial isolation and *in vitro* budding require approximately 6-7 hours, followed by a 16-hour ultracentrifugation step, tissue collection should be initiated early in the day.

### CRITICAL

All steps should be performed at 4 °C or on ice unless otherwise specified.

#### Innovation

- This protocol enables the isolation and analysis of distinct MDV subsets using density flotation gradients, allowing proteomic investigation of cargo selectivity across multiple MDV subsets.
- It provides a detailed workflow for MDV size determination and yield estimation by integrating transmission electron microscopy (TEM) with nanoparticle tracking analysis (NTA).
- The method is readily adaptable across mammalian systems and has been successfully applied to rat, rabbit, and bovine heart mitochondria.

#### Institutional permissions

All animal experiments were conducted following approval from the Institutional Animal Ethics Committee (IAEC), registered with the Committee for the Purpose of Control and Supervision of Experiments on Animals (CPCSEA), Government of India (Approval No. 326/GO/ReBiBt/S/2001/CPCSEA). All procedures, including modes of euthanasia (Wistar rats: CO_2_ inhalation; rabbits: overdose of thiopentone sodium), were carried out in accordance with national and institutional regulatory guidelines.

The protocol was optimized using rat heart tissue from adult animals (6–12 months old; ~250– 400 g; both sexes) and has also been successfully applied to rabbit heart tissue (adult animals >8 months; ~2–3 kg; both sexes).

For selected flotation experiments, bovine heart tissue was obtained from a local slaughterhouse. As this material was collected post-mortem from animals processed for commercial purposes, additional institutional animal ethics approval was not required.

#### Preparation of buffers and reagents

##### Timing: [one day before]

1. Prepare Mitochondria Isolation Buffer (MIB) (as per table below) and adjust the pH to 7.4, and store at 4 °C.
2. Prepare the buffer for budding reaction without succinate, ATP and ADP, which need to be added fresh.
3. Ensure antimycin A and N-ethylmaleimide (NEM) are reconstituted to final stock concentrations of 45.5 mM (25 mg/ml; 1000X stock in ethanol) and 100 mM (5X stock), respectively. The final concentrations of antimycin A and NEM in the *in vitro* budding reaction are 25 μg/ml and 20 mM, respectively.
4. Prepare an 80% (w/v) sucrose stock solution in cold mitochondrial isolation buffer (MIB). Dissolve sucrose completely at room temperature and store at 4 °C.
5. Prepare working sucrose solutions of 50%, 40%, 30%, 20%, and 10% by diluting the 80% sucrose stock with cold MIB. For gradients prepared in a SW41 Ti swinging-bucket rotor, 12 ml total gradient volume is required per tube (2 ml of each sucrose concentration). Keep all sucrose solutions on ice and prepare fresh on the day of use. **Note:** The gradient composition can be adjusted depending on experimental requirements. For example, a 30–60% sucrose gradient may be used to improve the separation of MDV subtypes, with 3 ml of each sucrose concentration per tube. For flotation gradients, the MDV-containing sample will be mixed with 80% sucrose to achieve a final concentration of 60% before loading at the bottom of the ultracentrifuge tube (see Sucrose Density Gradient Ultracentrifugation section). The ultracentrifugation rotor and adapters should be refrigerated before the experiment begins.

## Key resources table

| REAGENT or RESOURCE | SOURCE | IDENTIFIER |
|---|---|---|
| **Antibodies** |  |  |
| Pyruvate Dehydrogenase (C54G1) Rabbit mAb<br>(Used at 1:1000 dilution) | Cell Signaling Technology (CST) | 3205<br>RRID AB_2162926 |
| VDAC (D73D12) Rabbit mAb<br>(Used at 1:1000 dilution) | Cell Signaling Technology (CST) | 4661<br>RRID AB_10557420 |
| Purified Mouse Anti OPA1 (Used at 1:1000 dilution) | BD Transduction laboratories | 612606<br>RRID AB_399888 |
| TOM20 Rabbit mAb | Cell signaling technology | 42406<br>RRID AB_2799825 |
| Peroxidase-conjugated AffiniPure Goat Anti-Rabbit IgG (H+L) (Used at 1:5000 dilution) | Jacksons Immunoresearch Laboratories | 111-035-083<br>RRID AB_2313586 |
| Peroxidase-conjugated AffniPure Goat Anti-Mouse IgG (H+L) (Used at 1:5000 dilution) | Jacksons Immunoresearch Laboratories | 115-035-003<br>RRID AB_10015289 |
| **Chemicals, peptides, and recombinant proteins** |  |  |
| Mannitol | Himedia | MB198 |
| Sucrose | Himedia, Sigma | PCT0616, S0389 |
| Potassium chloride | Himedia | GRM3934 |
| Magnesium acetate | Himedia | MB039 |
| HEPES | Himedia | MB016 |
| ATP disodium salt | Himedia | RM439 |
| ADP disodium salt | Himedia | MB191 |
| Sodium succinate | Himedia | GRM418 |
| Dipotassium hydrogen phosphate | Himedia | TC596 |
| Sodium chloride | Himedia | GRM3954 |
| N-Ethylmaleimide | Sigma | E3876 |
| Antimycin A | Sigma | A8674 |
| GTP-γ-S | Sigma | G8634 |
| di-Sodium hydrogen phosphate dodecahydrate | Himedia | GRM1418 |
| Potassium dihydrogen orthophosphate monobasic | Himedia | GRM3943 |
| Dodecyl sulphate sodium salt | Himedia | TC698 |
| Tris buffer | Himedia | GRM1218 |
| Ammonium persulphate | Himedia | MB003 |
| TEMED | BIO RAD | 161-0800 |
| Glycine | Himedia | MB013 |
| Methanol | Thermo Fisher Qualigens | Q43637G |
| EGTA | Himedia | GRM1530 |
| Acrylamide | BIO RAD | 161-0101 |
| Bis-acrylamide | Himedia | MB005 |
| Tween-20 | Himedia | TC287 |
| Phosphotungstic acid hydrate | Sigma Aldrich | 79690 |
| EM grade gluteraldehyde | Merck | G5882 |
| Formvar/Carbon 200 Mesh, Gold grid | Electron Microscopy Sciences | FCF200-AU25 |
| **Experimental models: Organisms/strains** |  |  |
| Rat (Wistar) | RGCB animal facility |  |
| **Software and Algorithms** |  |  |
| Image J software | Schneider et al., 2012 | https://imagej.nih.gov/ij |
| Nanoparticle Tracking Analyser software (version 3.2) | Malvern Panalytical | n/a |
| **Other** |  |  |
| Sonicating water bath | Branson | 1800 |
| Shaking water bath | Julabo | SW22 |
| Ultracentrifuge | Beckman Coulter | L-100K |
| Swinging bucket Rotor | Beckman Coulter | SW41 Ti |
| Polyallomer centrifuge tubes | Beckman | 331372 |
| Nanoparticle tracking analysis | Malvern Panalytical | NanoSight 300NS |
| Transmission electron microscope | JEOL | JEM1101 |
| Transmission electron microscope | JEOL Ltd | JEM-2100 Plus |

### CRITICAL

Several chemicals listed above for downstream validation assays are hazardous and must be handled with care. For western blotting, acrylamide, bis-acrylamide, and TEMED are toxic. For transmission electron microscopy (TEM), glutaraldehyde and phosphotungstic acid are also hazardous. In particular, TEMED and glutaraldehyde can cause acute toxicity upon inhalation; therefore, they must be handled inside a certified chemical fume hood.

Appropriate personal protective equipment (PPE), including lab coat, safety goggles, and gloves, must be worn at all times, and all procedures should be performed in accordance with institutional safety guidelines.

## Materials and equipment

### Mitochondrial Isolation Buffer (MIB) (pH 7.4)

| **Component** | **Molecular weight** | **Final concentration** | **For 500 ml 2X MIB** |
|---|---|---|---|
| Mannitol | 182.7 Da | 220 mM | 40.194 g |
| Sucrose | 342.3 Da | 68 mM | 23.268 g |
| KCl | 74.55 Da | 80 mM | 5.964 g |
| EGTA | 380.35 Da | 0.5 mM | 190.175 mg |
| Magnesium acetate | 214.45 Da | 2 mM | 428 mg |
| HEPES pH 7.4 | 238.3 Da | 20 mM | 2.383 g |
| Milli-Q | n/a | n/a | Make up to 500 ml |

Note: Store the 2X buffer at 4 <sup>o</sup>C. We have had no reproducibility issues up to 6 months of storage.

### Budding Reaction Buffer (BR) (pH 7.4)

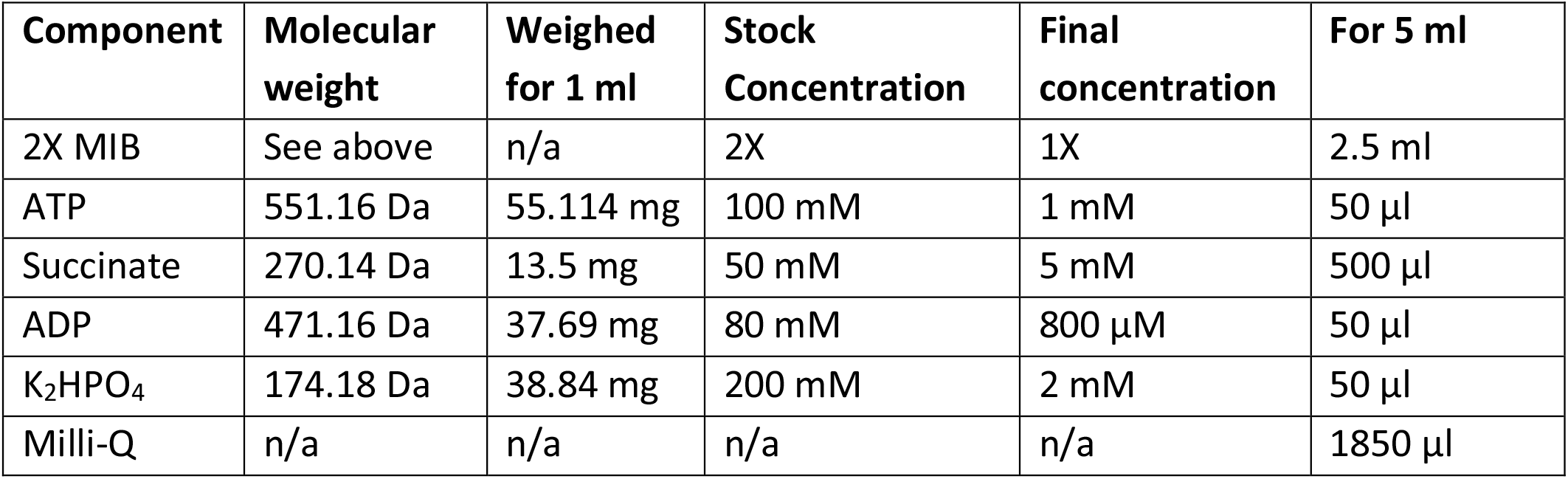

| **Component** | **Molecular weight** | **Weighed for 1 ml** | **Stock Concentration** | **Final concentration** | **For 5 ml** |
|---|---|---|---|---|---|
| 2X MIB | See above | n/a | 2X | 1X | 2.5 ml |
| ATP | 551.16 Da | 55.114 mg | 100 mM | 1 mM | 50 µl |
| Succinate | 270.14 Da | 13.5 mg | 50 mM | 5 mM | 500 µl |
| ADP | 471.16 Da | 37.69 mg | 80 mM | 800 µM | 50 µl |
| K<sub>2</sub>HPO<sub>4</sub> | 174.18 Da | 38.84 mg | 200 mM | 2 mM | 50 µl |
| Milli-Q | n/a | n/a | n/a | n/a | 1850 µl |

| **Component** | **Molecular weight** | **Final concentration** | **For 1 L** |
|---|---|---|---|
| NaCl | 58.4 Da | 137 mM | 8 g |
| KCl | 74.55 Da | 2.7 mM | 0.2 g |
| Na<sub>2</sub>HPO<sub>4</sub> | 358.14 Da | 10 mM | 1.44 g |
| KH<sub>2</sub>PO<sub>4</sub> | 136.09 Da | 1.8 mM | 245 mg |
| Milli-Q | n/a | n/a | n/a |

## Step-by-step method details

### Mitochondrial Isolation from Rat Heart

#### Timing: [3 hours]

1. Sacrifice animals in accordance with approved institutional guidelines. We have sacrificed rats using CO_2_ inhalation. Excise heart and immediately transfer into ice-cold phosphate-buffered saline (PBS).
2. Wash the hearts 2-3 times in ice-cold PBS to remove excess blood (Figure 1A).
3. Using sterile scalpel and forceps, chop the tissue into ~5 mm^3^ pieces in a Petri dish containing ice-cold PBS (Figure 1B).
4. Transfer the chopped tissue into a pre-chilled 50 ml conical tube and centrifuge at 3,000 g for 10 min at 4 °C. **Note:** All centrifugation steps for mitochondrial isolation were done using an Eppendorf 5810R bench-top centrifuge with a fixed-angle rotor (F-34-6-38). Discard the supernatant, and wash the tissue pellet three times using 10 ml ice-cold PBS at 3,000 g at 4 °C for 10 min (Figure 1C).
5. Homogenize the pellet using a Dounce homogenizer (~80 times) into 10 ml of cold mitochondrial isolation buffer (Figure 1D). We have used a manual method with the 50 ml homogenizer with a type A (loose) pestle. For bovine heart, semi-automatic homogenization using Schuett homgen^plus^ homogenizer was used.
6. Centrifuge the homogenate at 3,000 g for 10 min at 4 °C to pellet unbroken cells, nuclei, and debris. Carefully transfer the supernatant to a new pre-chilled tube without disturbing the pellet.
7. Repeat the low-speed centrifugation step two additional times, each time transferring the supernatant to a new pre-chilled tube. This step improves the removal of contaminating nuclei and debris
8. Centrifuge the supernatant from the last step at 10,000 g at 4 °C for 20 min. The pellet is enriched in heavy membranes, including mitochondria (Figure 1E). **Note:** The post-mitochondrial supernatant from this step is retained for cytosol preparation. To obtain cytosol, ultracentrifuge the supernatant at 200,000 g for 90 min at 4 °C to remove residual membranes. The resulting supernatant represents the cytosolic fraction and can be used for *in vitro* budding reactions.
9. Resuspend the mitochondrial pellet in 5 ml cold MIB and wash the mitochondrial pellet by centrifuging at 10,000 g at 4 °C for 10 min.
10. Resuspend the final pellet in 1 ml budding reaction buffer and transfer it into a fresh prechilled 1 ml microfuge tube. Collect 5 μl of the sample to determine concentration using the Bradford assay and assess mitochondrial yield.

**Figure 1:**
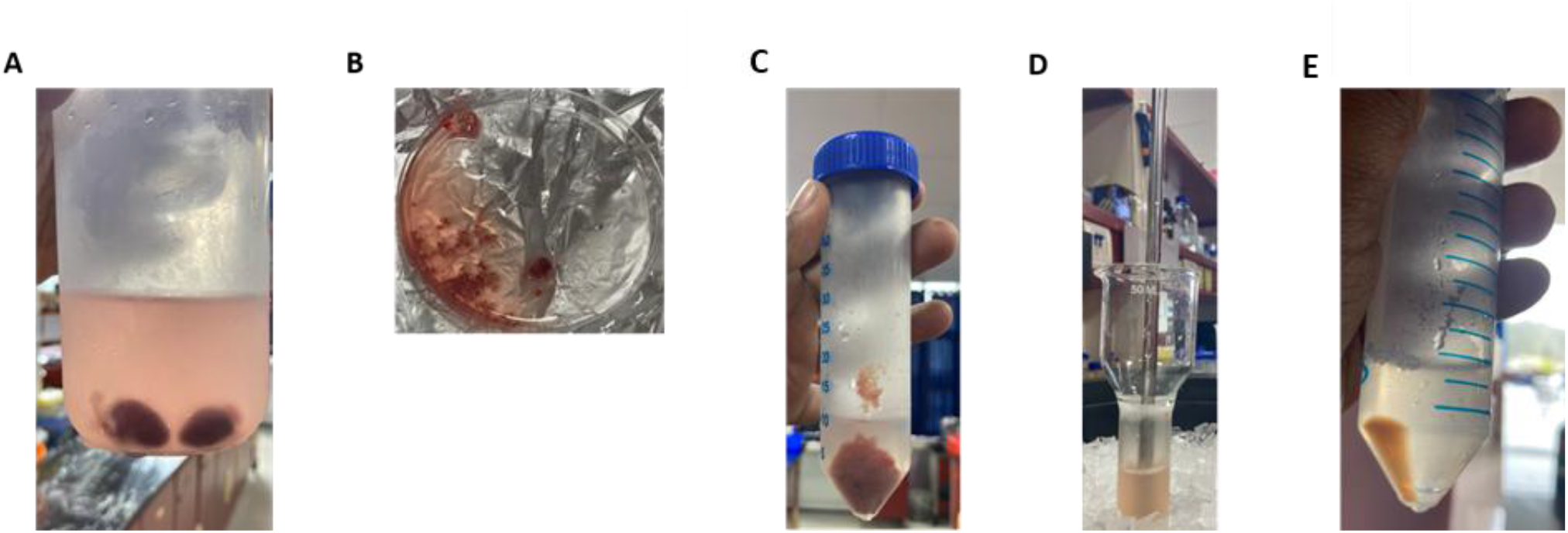
Stepwise isolation of mitochondria from rat heart tissue. (A) Rat heart in PBS (B) Tissue after mincing with a scalpel (C) Pellet before homogenisation (D) Dounce homogenisation (E) Mitochondria-enriched final pellet yield from 2 rat hearts after 10,000 g centrifugation.

### *In Vitro* Budding of MDVs

#### Timing: [3 hours]

Here, we describe the in vitro budding reaction with two major controls: antimycin A treatment of mitochondria, which increases ROS generation by inhibiting complex III, and NEM, an alkylating agent that inhibits *in vitro* trafficking and is reported to inhibit MDV generation ^[2]^.

11. Set up budding reactions under the following conditions: control and NEM (20 mM) and Antimycin A (50 μM). Reactions are prepared in a budding reaction buffer supplemented with ATP, ADP, and succinate at the final concentrations specified in the table above. For each condition, use 2-5 mg of the isolated mitochondria in a total reaction volume of 500 μl. **Note:** We make a 2X MIB stock buffer so that the addition of ATP, ADP, succinate and K_2_HPO_4_ and the addition of NEM, does not dilute the final budding reaction buffer. We make up the reaction with water (total 500 μl) to achieve the final 1X concentrations. We add the substrate and inhibitors simultaneously to the mitochondria.
12. Incubate the reaction for 2 hours at 37 °C in a water bath with mild intermittent sonication or shaking at 80 rpm. **Note:** Ensure the temperature does not exceed 37 °C during the reaction, as sonication can raise the water bath temperature. To minimize heating, sonicate for 10–15 seconds at 10-minute intervals (Branson, 1800 Series; 40kHz). Alternatively, incubation in a shaking water bath (Julabo SW22) at 37 °C and 80 rpm can be used to promote budding. In our hands, these dynamic conditions yielded improved MDV recovery compared with static water-bath incubations (see Expected Outcomes).
13. Centrifuge the sample at 10,000 g at 4 °C for 10 min to remove intact mitochondria.
14. The supernatant containing MDVs is again centrifuged 3-4 times at 10,000 g until there is no visible mitochondrial pellet.
15. The MDVs present in the supernatant are subjected to flotation density gradient ultracentrifugation to further enrich for MDVs. **Note:** Isolated MDVs can be stored for four days in 4 °C **(See Troubleshooting Problem 5)** or used for downstream analyses after protein quantification **(See Troubleshooting Problem 1)**.

### Sucrose Density Gradient Ultracentrifugation

#### Timing: [16 hours]

16. Mix 500 μl of the MDV sample from the last step with 1.5 ml of 80% sucrose prepared in MIB to achieve a final sucrose concentration of 60%. Load the sample at the bottom of the ultracentrifuge tube.
17. Sequentially overlay the sample with 2 ml each of 50%, 40%, 30%, 20%, and 10% sucrose solutions, forming a discontinuous gradient. When layering each sucrose solution, slowly dispense the solution along the inner wall of the tube, positioning the pipette tip slightly above the interface of the previous layer. Add the solution gently to avoid disturbing the underlying gradient and maintain sharp layer interfaces (Figure 2A, B). **Note:** The polyallomer ultracentrifugation tubes have a volume of 13 ml (SW41 Ti swinging bucket rotor). The topmost 1 ml is left free, and the total volume is 12 ml.
18. Centrifuge the gradients at 200,000 g for 16 h at 4 °C.
19. Following centrifugation, carefully collect 1 ml fractions from the top of the gradient (Figure 2C, D), yielding 12 fractions for downstream processes, including western blot analysis for MDV cargo. **(See Troubleshooting Problem 2)**

**Figure 2:**
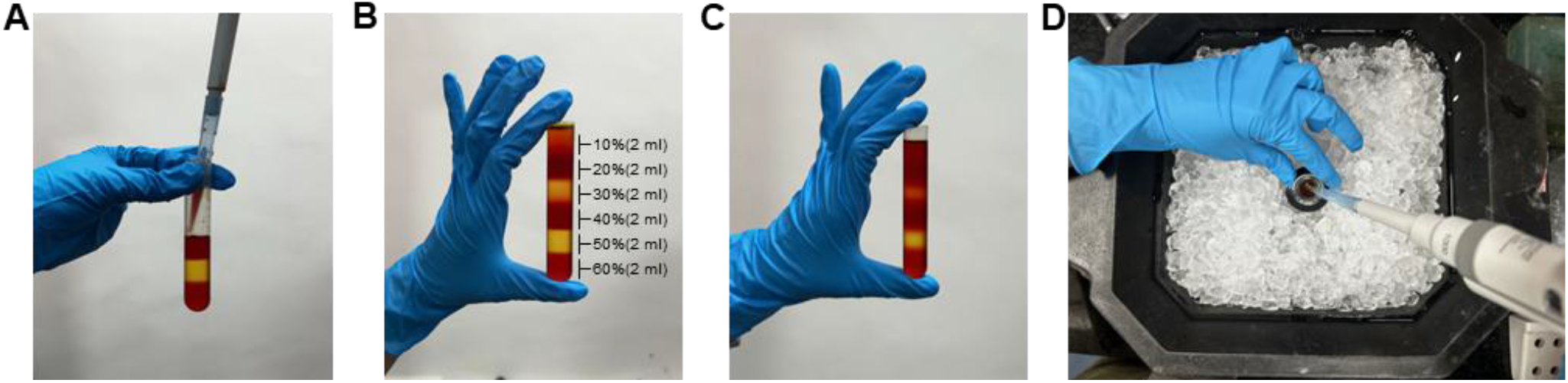
Preparation and fractionation of discontinuous sucrose gradients for MDV isolation. (A) Representative image of the discontinuous sucrose gradient layering process in an ultracentrifuge tube. We carefully overlay sucrose solutions (10–50%) onto the 60% sucrose cushion by slowly dispensing each layer along the inner wall of the tube, slightly above the previous interface to preserve gradient integrity. For visualization, we add food coloring to the sucrose solutions to enhance contrast between layers; this is not used in the experiment. (B) Gradients prior to ultracentrifugation, with 2 ml of each sucrose layer sequentially overlaid. We place tubes into swinging-bucket holders and precisely balance them by adjusting the weight with MIB buffer added to the top of the gradient. (C) Gradients following ultracentrifugation at 200,000 g (35,000 rpm in an SW41 Ti rotor), showing slight intermixing at interfaces after centrifugation. (D) Fraction collection from the top of the gradient. We carefully collect fractions on ice to preserve vesicle integrity.

#### Pause point

Please note that until the sample is kept overnight for flotation ultracentrifugation, there are no pause points. The mitochondrial isolation must be followed immediately by the budding reaction.

### Nanoparticle tracking analysis (NTA) of density-floated MDVs

Nanoparticle tracking analysis (NTA) is performed to determine MDV size distribution and concentration. NTA measurements are performed using a NanoSight NS300 system (Malvern Panalytical). Sucrose present in density gradient fractions can interfere with NTA by increasing solution viscosity and affecting particle tracking **(See Troubleshooting Problem 3)**. Therefore, MDV fractions obtained from sucrose gradients are diluted prior to NTA analysis in the following steps.

20. Dilute 30% and 50% sucrose fractions in 1X MIB lacking sucrose and mannitol to reduce the final sucrose concentration to <10%. This is achieved by a 3-fold dilution of the lighter interface and a 5-fold dilution of the heavier interface.
21. Concentrate the diluted fractions by centrifugation at 14,000 g using 3 kDa Amicon centrifugal filter units to restore vesicle concentration.
22. Switch on the NTA instrument and allow it to stabilise for 10-15 min. Thoroughly clean the sample chamber with distilled water and ensure there are no air bubbles.
23. Prepare a blank (negative control) using 1 ml of filtered PBS.
24. Load the blank into the chamber slowly and confirm the absence of detectable particles before proceeding with the sample.
25. Dilute samples in filtered 1X PBS (typically 10:990 to 100:900) to obtain an optimal particle concentration of 20–100 particles per frame. Ensure that the final sucrose concentration remains below 5% **(see Troubleshooting Problem 3**).
26. Capture particle movement videos (10s) using CMOS acquisition mode with the following settings: 300-400 frames per recording and slider shutter and slider gain set to 1300 and 512, respectively (see Video S1 for a representative video).
27. Record each sample at least five times for reproducibility. Avoid bulk-flow artefacts and ensure at least 1000 completed tracks are obtained. Optimal measurement accuracy is typically achieved at particle concentrations of 10^7^-10^9^ particles/ml. This may vary for each NTA instrument; please check the optimal concentrations for the brand of instrument used.
28. Analyze data using Nanoparticle Tracking Analyser software (version 3.2) to obtain vesicle size distribution and particle concentration (See Figure 4 for concentration vs size plot of total budding reaction, heavy and light MDVs). **Note:** Following NTA, the MDV ultrastructure is analyzed by TEM and its cargo proteomically determined. **(See Troubleshooting Problem 4)**

## Expected outcomes

For rat hearts, we typically obtain ~10 mg of mitochondria per rat heart and generally pool 2-3 hearts to set up reactions in technical duplicates. Following the in vitro budding reaction, the post-budding supernatant contains ~15-25 µg protein under control conditions, which decreases to <10 µg upon addition of N-ethylmaleimide (NEM). After sucrose gradient fractionation, protein levels fall below the detection limit of the Bradford assay. We interpret the residual protein detected after NEM treatment as non-vesicular contaminating material rather than bona fide MDVs. Control and NEM-treated budding reactions were set up with equal amounts of mitochondria, and equal volumes of control and NEM-treated budding reactions were analysed (Figure 3A). Under these conditions, nearly all previously reported MDV cargo proteins are reduced to near-undetectable levels in the MDV fraction upon NEM exposure. NEM is a known thiol-alkylating agent and hence inhibits cytosolic trafficking factors that require free thiol groups for vesicle formation. The marked depletion of mitochondrial cargo proteins in the MDV fraction supports the conclusion that their presence in control reactions arises from an active budding process. In our assay, we therefore define NEM sensitivity as the specific reduction or loss of mitochondrial protein signal in the MDV fraction upon NEM treatment, and use this criterion to distinguish true MDV cargo from non-specific protein release due to loss of membrane integrity. This also serves as the basis for our subtraction proteomics of the budding reaction^[1]^.

**Figure 3:**
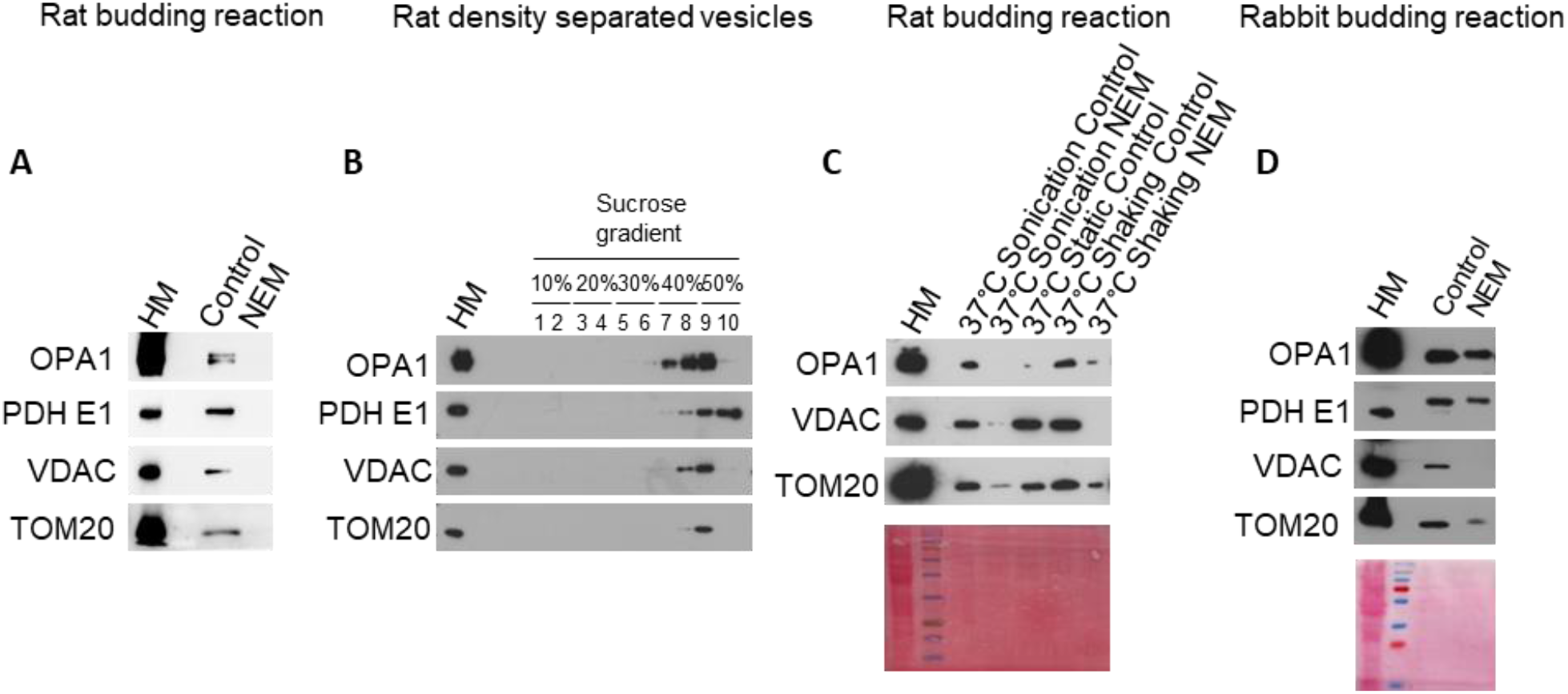
Biochemical validation of *in vitro* generated vesicles. (A) Western blot showing NEM sensitivity of rat cardiac MDV cargo. OPA1, PDH E1, VDAC, and TOM20 are probed in MDV fractions generated in the presence or absence of N-ethylmaleimide (NEM). (B) Density flotation analysis of MDVs in a 10-60% discontinuous sucrose gradient. Marker proteins, including OPA1 (inner membrane/intermembrane space), pyruvate dehydrogenase E1 (PDH E1; matrix), and VDAC (outer membrane), are probed across sucrose fractions. (C) Effect of incubation conditions on MDV yield. Shaking and mild sonication enhance MDV recovery compared to static incubation. Vesicles generated under these dynamic conditions remain NEM-sensitive, demonstrating that increased yield reflects enhanced active budding rather than nonspecific membrane fragmentation. (D) Western blot showing NEM sensitivity of rabbit cardiac MDV cargo. OPA1, PDH E1, VDAC, and TOM20 are probed in MDV fractions generated in the presence or absence of N-ethylmaleimide (NEM).

We have observed that budding reactions can be initiated with as little as 0.5 mg of mitochondria and still yield a western detectable mitochondrial protein signal in the MDV fraction (data not shown). However, for density flotation experiments, budding reactions are set up using a minimum of 5 mg mitochondria to ensure sufficient material for fractionation and downstream analyses. Following 10-60% sucrose flotation ultracentrifugation, most MDVs peak at fraction 9. Vesicles containing OPA1 display a broader density distribution, suggesting incorporation into multiple MDV subsets, whereas Pyruvate dehydrogenase E1 (PDH E1), a matrix protein, is enriched in higher-density fractions (Figure 3B), consistent with the presence of biochemically distinct vesicle subsets. To further resolve distinct MDV subsets, a 30-60% sucrose gradient can be used. Under these conditions, higher-density, matrix-containing vesicles are more clearly separated from lower-density OPA1-containing MDVs, facilitating improved discrimination of MDV subtypes.

MDV formation is temperature-dependent. When budding reactions are performed under shaking conditions at 37 °C and 4 °C, incubation at 4 °C results in a marked reduction of all proteins examined in the MDV fraction (refer Rajeev T et al)^[1]^. This temperature dependence is consistent with previous *in vitro* MDV budding studies^[2]^ and validates 4 °C as an appropriate negative control, supporting the conclusion that MDV formation is an active process. In addition, shaking and mild sonication yield higher MDV yields than static incubation. Importantly, vesicles generated under shaking or sonication conditions remain NEM-sensitive, with a marked reduction of mitochondrial cargo in the MDV fraction upon NEM treatment (Figure 3C). The NEM sensitivity of the cargo supports an active budding process.

In reactions supplemented with cytosol (2 µg/µl), we included both NEM and GTPγS (0.1 mM) controls and observed that with cytosol, as expected, there was an increased incorporation of OPA1 and PDH E1. However, addition of GTPγS increased recovery of OPA1 but not PDH E1 in the MDV fraction (refer Rajeev T et al)^[1]^. This is in line with a previous study showing that the addition of GTPγS increased the incorporation of VDAC into the MDV fraction^[2]^. This suggests that in the *in vitro* context, vesicle scission may occur without Drp1-mediated GTP hydrolysis. Some GTPases can become locked in an active state with GTPγS, which may drive better cargo selection of OPA1.

We extended the assay to rabbit heart mitochondria and observed similar MDV budding behavior as in rat heart mitochondria (Figure 3D). MDV cargo incorporation was assessed by probing marker proteins across the budding reaction. In our hands, OPA1, PDH E1, VDAC, and TOM20 are readily detected in the MDV fraction and show clear sensitivity to NEM, consistent with inhibition of vesicle formation under these conditions.

Our overall aim was to establish a robust MDV isolation protocol suitable for downstream applications such as transmission electron microscopy (TEM), NTA and proteomics. Prior to ultrastructural and proteomic characterization, we performed NTA to assess vesicle size distribution and particle concentration during budding reactions, confirming the presence of vesicles within the expected nanoscale range (Figure 4A). The heavy fraction showed a slightly smaller median diameter than the light fraction, but this difference was small. Transmission electron microscopy of budding reactions reveals both single- and double-membrane vesicles. A clear reduction in vesicular structures is observed after NEM treatment (Figure 4B). TEM of the heavy and light MDV fractions recovered by density flotation shows vesicular structures in both fractions (Figure 4C).

**Figure 4:**
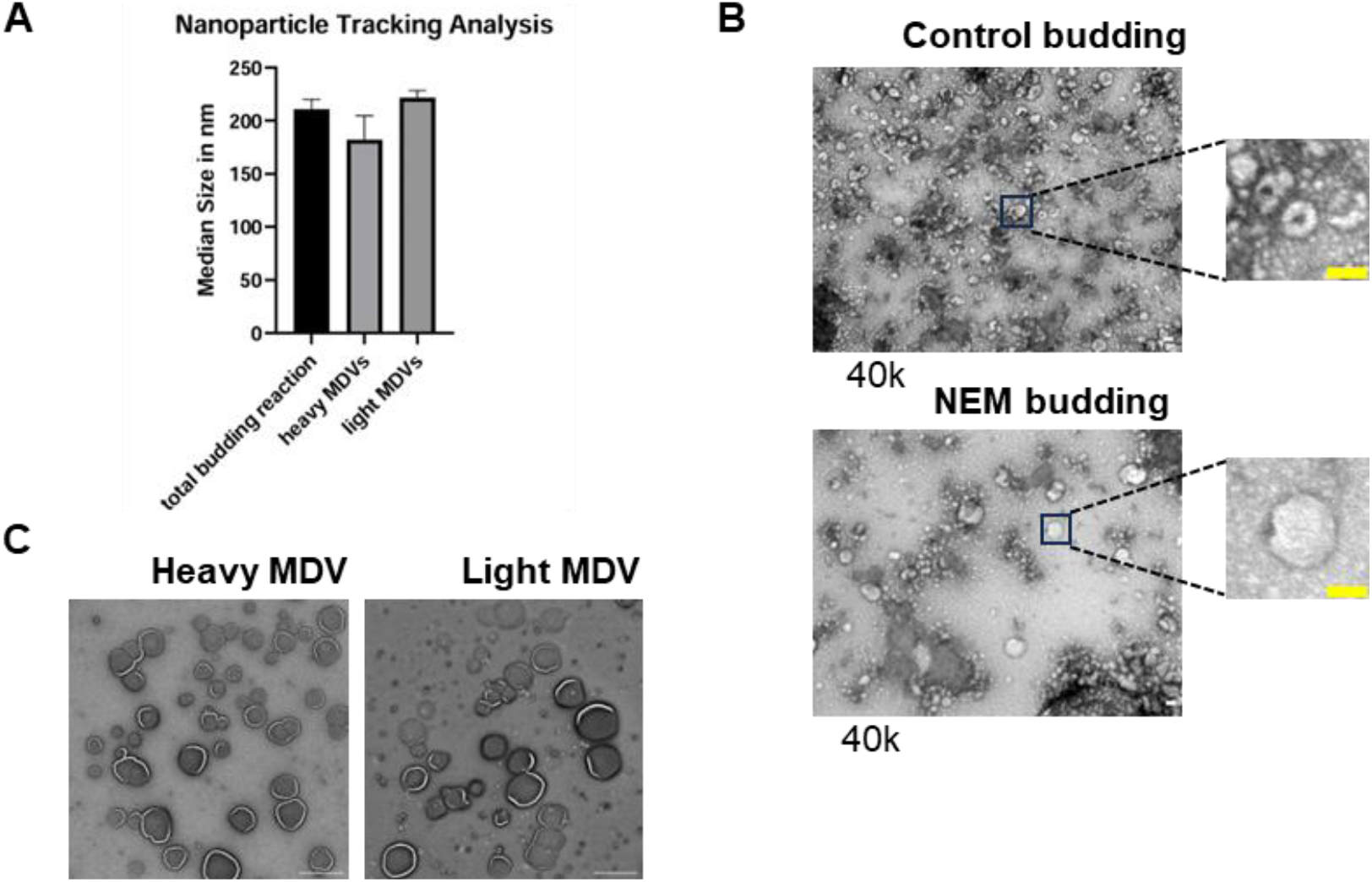
Quantitative and ultrastructural validation of *in vitro* generated MDVs. (A) Nanoparticle tracking analysis (NTA) of budding reactions and floated MDVs. (B) Transmission electron microscopy (TEM) of MDVs generated without and with N-ethylmaleimide (NEM) (median diameter 85 nm). A marked reduction in vesicle number is observed following NEM treatment. Scale bar = 100 nm. (C) Transmission electron microscopy (TEM) of density-purified MDVs. 50-60% interphase fractions constitute the heavy MDVs and 30-40% interphase fractions constitute the light MDVs. Scale bar = 1 µm.

**Figure 5:**
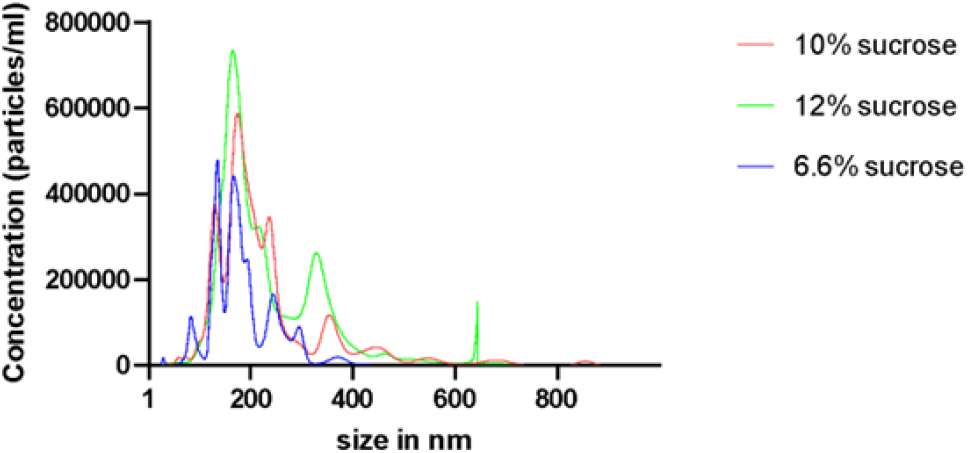
NTA of sucrose solutions ranging from 6-12%.

Following proteomic analysis of density-floated MDVs, approximately 28-35% of the detected proteins are of mitochondrial origin based on the validated MitoCarta 3.0 reference list. This is comparable to previously reported MDV proteomic dataset of rat cardiac-derived MDVs at 35%-40% sucrose interface, which identified 250 mitochondrial proteins among 714 total proteins (~35%), ^[3]^. Together, these comparisons suggest that density flotation coupled with an NEM sensitivity criterion identifies true MDV cargo for downstream analyses.

Overall, this protocol yields NEM-sensitive, temperature-dependent budded MDVs that can be quantitatively assessed, ultrastructurally validated, and proteomically characterized, providing a robust framework to investigate MDV biogenesis and cargo selectivity.

## Limitations

A key limitation of this protocol is the requirement for large amounts of mitochondria, which are difficult to obtain from cultured cells. As a result, rapid cell culture–based experiments to study MDV biogenesis are not feasible. An alternative method for obtaining an MDV proteome within cells was reported by the Heidi McBride group ^[4]^. They generate the TOM20^+^ MDV proteome by dual-tagging the OMM proteins MAPL and TOM20 in COS7 cells, and since these are independently packaged into MDVs, TOM20^+^ MDVs could be separated from dual-positive OMM fragments ^[4]^. This represents a steady-state MDV proteome from cells. However, MDVs generated *in vitro*, as described here, complement the previous approach, particularly for MDVs from common model organisms such as rats, mice, and rabbits, which are particularly useful for cardiovascular research. They would be useful for reconstitution approaches to understand the biogenesis and trafficking steps in greater detail.

It should be noted that Antimycin A induced MDVs do not consistently show increased cargo incorporation for all MDV cargo. We observed that while OPA1 levels increased with Antimycin treatment, VDAC levels decrease under the same conditions (data not shown). Previous study^[2]^ had observed that VDAC was similarly less incorporated into MDVs upon Antimycin treatment. While comparing the proteomes of control MDVs to Antimycin A-treated MDVs previous studies observed almost all proteins in MDVs did not show an increase after antimycin A treatment^[3, 5]^. This reductionist approach where the mitochondria is removed from the cellular environment including loss of ER-mito contact sites may therefore fail to recapitulate budding under oxidative stress. However, as our approach resolves multiple sub-types of basal MDVs, it can still be a powerful tool to understand how biogenesis of even the energetically demanding double-membrane vesicles occur.

## Troubleshooting

### Problem 1

#### Low MDV yield despite adequate mitochondrial input (related to step 15)

##### Potential solutions

- Assess mitochondrial integrity prior to budding and after budding (TMRE staining to assess mitochondrial potential).
- Ensure succinate, ATP, and ADP are freshly added to the budding buffer immediately prior to use.
- Set up the budding reaction immediately after mitochondrial isolation to preserve mitochondrial integrity and functionality.

### Problem 2

#### No signal for proteins in Western Blots after flotation ultracentrifugation (related to step 19)

##### Potential solutions

- Always include a mitochondrial pellet control to verify antibody performance.
- Increase the amount of mitochondria used for the budding reaction. Although budding can be initiated with as little as 0.5 mg mitochondria, a western detectable signal in density flotation experiments requires a minimum of 5 mg mitochondria to ensure sufficient material for fractionation and downstream analyses (related to step 10).
- If mitochondrial yield from rat heart preparations is limiting (~10 mg per heart), consider scaling up the number of hearts pooled per experiment.
- For scaling-up flotation further, bovine heart mitochondria may be used as an alternative source. From ~40 g of bovine heart tissue, 20–25 mg mitochondria can typically be obtained, enabling larger-scale budding reactions.

### Problem 3

#### Estimating yield and size range of MDVs

Nanoparticle tracking analysis (NTA) tends to overestimate vesicle sizes as it has reduced sensitivity for small/low-scattering particles. It can be biased toward detecting brighter, slower-moving particles, which have to be cross-confirmed with TEM. In our hands, using this protocol, vesicle size distribution measured by NTA has a median around 200 nm and TEM shows a median of 85 nm confirming previous MDV literature. This suggests that indeed we might miss out smaller vesicles that don’t scatter well in the NTA analysis. Additionally, sucrose in the density-floated fractions can interfere NTA (related to step 25).

##### Potential solutions

- Analyze freshly prepared post-budded MDVs whenever possible.
- We performed NTA on sucrose solutions ranging from 6 to 12% and observed particles with sizes comparable and smaller than MDVs. However, the concentrations were very low in the range of 10^5^-10^6^ particles, <250 completed tracks and almost 100 to 1000-fold lower than our vesicle samples. To minimize potential interference from sucrose during measurement, NTA of density-floated MDV samples was performed only after diluting the fractions to reduce the sucrose concentration to below 5%. In our hands, maintaining the final sucrose concentration below 1% yields reliable particle detection and size estimation.
- Include appropriate negative controls (e.g., NEM-treated reactions or buffer-only controls) to distinguish *bona fide* MDVs from background particles or sucrose-related impurities/artefacts.

### Problem 4

#### High background or non-mitochondrial contamination in proteomics (related to step 28)

##### Potential solutions

- Use NEM sensitivity as a criterion to define *bona fide* MDVs. Perform parallel budding reactions in the presence and absence of NEM and apply a subtractive proteomics strategy, prioritizing proteins that are reduced or lost upon NEM treatment. We have successfully implemented this NEM-based subtractive proteomics approach to define MDV-enriched cargo^[1]^.
- MDVs are heterogeneous, with different cargo compositions and potentially distinct biogenesis pathways. Sucrose density flotation of isolated MDVs allows characterization of distinct vesicle populations, providing insight into subtype-specific cargo selection and functional differences.
- Treat isolated MDVs with trypsin followed by proteomic analysis of the trypsin-protected fraction. This distinguishes true MDV cargo from non-vesicular mitochondrial proteins.

### Problem 5

#### Sample storage (related to step 15)

##### Potential solution

We typically proceed directly from the budding reaction to density gradient flotation on the same day without intermediate storage. However, MDV samples stored at 4 °C for up to 4 days prior to adsorption onto TEM grids did not show detectable changes in vesicle size distribution or morphology.

### Resource availability

#### Lead contact

Further information and requests for resources and reagents should be directed to and will be fulfilled by the lead contact, Dr. Ananthalakshmy Sundararaman.

#### Technical contact

Technical questions on executing this protocol should be directed to and will be answered by the technical contact, Thejaswitha Rajeev

#### Materials availability

This study did not generate new unique reagents.

## Data and code availability

The study did not generate datasets apart from the ones presented. No codes were used for analysis. Some elements of the graphical abstract were created in Biorender.

## Acknowledgments

We acknowledge DBT Ramalingaswami grant to AS and the UGC fellowship to TR for funding. The TEM facility, animal facility and central ultracentrifugation facility of RGCB played a key role in this study. We also acknowledge the Central Instrumentation Facility at CSIR-IICB for assistance with NTA analysis.

## Author contributions

1. Thejaswitha Rajeev: Rat and Rabbit heart MDV generation and proteomics, standardisation of parameters for vesicle budding, flotation-based purification of vesicles.
2. Ela Lepcha: Nanoparticle tracking analysis of MDVs
3. Nidhi Nair: Standardisation of parameters for vesicle budding and flotation-based purification of vesicles
4. Lariza Ramesh: Rat and Rabbit heart MDV generation and independent validation of the protocol.
5. Areeba Marib: Standardisation of parameters for mitochondrial isolation and budding.
6. John B Johnson: Mounting and negative staining of vesicles, TEM imaging and analysis
7. Dhabaleswar Patra: Nanoparticle tracking analysis of MDVs
8. Ananthalakshmy Sundararaman: conceived and coordinated the study and wrote the paper.

All authors analysed the results and approved the final version of the manuscript.

## Declaration of interests

The authors declare no competing interests.

## Notes

### Competing Interest Statement

The authors have declared no competing interest.

### Summary of Updates

An additional section on NTA analysis of MDVs, along with additional authors and their affiliations, has been added to the revised manuscript.

